# elPrep 4: A multithreaded framework for sequence analysis

**DOI:** 10.1101/492249

**Authors:** Charlotte Herzeel, Pascal Costanza, Dries Decap, Jan Fostier, Wilfried Verachtert

**Affiliations:** ExaScience Life Lab, imec, Leuven, Belgium; Department of Information Technology, Ghent University - imec, Ghent, Belgium

## Abstract

We present elPrep 4, a reimplementation from scratch of the elPrep framework for processing sequence alignment map files in the Go programming language. elPrep 4 includes multiple new features allowing us to process all of the preparation steps defined by the GATK Best Practice pipelines for variant calling. This includes new and improved functionality for sorting, (optical) duplicate marking, base quality score recalibration, BED and VCF parsing, and various filtering options. The implementations of these options in elPrep 4 faithfully reproduce the outcomes of their counterparts in GATK 4, SAMtools, and Picard, even though the underlying algorithms are redesigned to take advantage of elPrep’s parallel execution framework to vastly improve the runtime and resource use compared to these tools. Our benchmarks show that elPrep executes the preparation steps of the GATK Best Practices up to 13x faster on WES data, and up to 7.4x faster for WGS data compared to running the same pipeline with GATK 4, while utilizing fewer compute resources.

## Introduction

elPrep 4 is a vastly extended reimplementation of elPrep [1], a multithreaded tool for preparing sequence alignment/map files (SAM/BAM) [2] for variant calling in DNA sequencing pipelines. Which preparation steps are used in a pipeline depends on the application, but, in general, they prepare the aligned read data in some way for statistical analysis, and they may include steps for filtering out unmapped reads or reads based on genomic regions of interest, sorting reads for coordinate order, marking the reads that are optical or PCR duplicates, calculating and applying base quality score recalibration, and so on. The GATK Best Practices [3] for example define a 4-step pipeline –and a couple of variations– for preparing data for variant calling with GATK [4], one of the most widely used variant callers.

elPrep differs from other tools for processing SAM/BAM files such as SAMtools [5], Picard, and GATK 4 [4]^1^ in terms of its software architecture that allows executing sequencing pipelines by making only a single pass through the data, independent of the number of steps specified in the pipeline description. This software architecture is designed to avoid repeated file I/O by keeping data as long as possible in memory during execution, to merge the computations of different preparation steps, and to avoid unnecessary synchronization while parallelizing execution, all of which significantly reduce the time needed to execute a sequencing pipeline [1].

elPrep 4 is a complete redesign and reimplementation of elPrep [1] in Go, an open-source programming language developed by Google. Go is a statically typed, compiled language featuring memory safety, parallel garbage collection, type inferencing, and support for concurrency utilizing multiple cores, which gives us access to new software optimization strategies to further improve the performance of elPrep. The original implementation of elPrep was written in Common Lisp, a language with good support for low-level performance optimizations thanks to optional type declarations, code inlining, stack-based memory allocation, and multithreading features.

One aspect specific to a sequencing application such as elPrep is that it needs to process hundreds of gigabytes of data, putting a tremendous pressure on memory management [6]. Common Lisp uses a stop-and-copy, stop-the-world garbage collector, which we needed to turn off because it interfered too much with the multithreaded execution of elPrep as it frequently pauses the program. Without garbage collection, we needed to employ a rigid programming style where we reuse memory and avoid unnecessary memory allocation as much as possible, increasing the complexity for programming and maintaining elPrep. Go comes with a concurrent, parallel garbage collector which solves this problem [6]. Other advantages of switching to Go include its portable, free compiler and modern language features such as type inferencing, UTF8 by default, escape analysis by the compiler, and so on.

The new elPrep 4 framework also allows us to more easily add new functionalities, and to implement all of the preparation steps described by the GATK Best Practices [3]. Two key contributions include algorithms for optical duplicate marking and base quality score recalibration, both optimized for efficient parallel execution in the elPrep framework, while producing the same results compared to their respective implementations in Picard and GATK 4. This involves a non-trivial reformulation of these algorithms that, compared to the original algorithms in Picard and GATK 4, avoid the use of intermediate files, avoid multiple iteration loops over the data, and are parallel.

We show that elPrep 4 drastically reduces the runtime and resource cost for running sequencing pipelines by benchmarking a 4-step pipeline from the GATK Best Practices in elPrep and comparing it to both the GATK 3.8 and GATK 4 runtimes. We also discuss a scaling experiment on Amazon Web Services (AWS) that compares the dollar cost of running elPrep 4 versus GATK 4 to process both whole-exome and whole-genome data.

## Implementation

elPrep is developed at the ExaScience Life Lab (http://www.exascience.com) for the Linux operating system. elPrep 4 is written in Go, a programming language developed by Google. Source code and documentation are available at http://github.com/ExaScience/elprep under the terms of the GNU Affero General Public License version 3 as published by the Free Software Foundation, with Additional Terms. Demos and test data can be downloaded from our Github repository at http://github.com/ExaScience/elprep/tree/master/demo.

## Materials and methods

elPrep 4 extends and improves on the original elPrep [1] functionality. For example, with elPrep 4 it is possible to execute all preparation steps recommended by the GATK Best Practices [3] for variant calling, but it can also be used for implementing other types of pipelines [7]. We present an overview of the newly added functionality, as well as the non-trivial algorithms we designed to implement this.

### elPrep 4 overview

elPrep 4 introduces the following new features:

1. Base quality score recalibration (BQSR): We added an option (–bqsr) to perform BQSR. This option essentially combines the semantics of the GATK 4 commands BaseRecalibrator and ApplyBQSR, producing identical results.
2. Optical duplicate marking: We added an option (–mark-optical-duplicates) to perform optical duplicate marking. The Picard/GATK 4 option for duplicate marking (MarkDuplicates) automatically performs optical duplicate marking after a generic duplicate marking phase based on adapted mapping positions of reads. The optical duplicate marking phase is used to generate metrics to distinguish between PCR and optical duplicates. The –mark-optical-duplicates option tells elPrep 4 to do the same.
3. Metrics: elPrep now generates metrics files that contain statistics about the number of unmapped reads, secondary reads, read duplicates, base quality scores, etc. It has the option to output the same metrics as the .metrics and .recal metrics generated by Picard/GATK 4. The format of the elPrep metrics files is identical to those from Picard/GATK 4 and are compatible with MultiQC [8] for visualization.
4. BAM parsing: elPrep 4 previously relied on calling SAMtools for BAM parsing, but now implements BAM parsing itself using the built-in gzip compression library of Go. The compression is now more efficient in terms of runtime.
5. VCF parsing: elPrep 4 provides VCF parsing. This was implemented to handle the known sites (cf. dbsnp files) for base quality score recalibration, but can be used to implement other tools.
6. Filtering reads based on genomic regions specified by a BED file: This is an option similar to the -L options in SAMtools/Picard/GATK. We added BED file parsing to elPrep to support this.
7. Integrated split-filter-merge (sfm) mode: elPrep offers two execution modes, namely a mode that operates entirely in RAM, and a mode that splits data using genomic regions for processing (sfm). This was previously implemented using Python scripts, but these are now replaced by an sfm subcommand implemented in Go as well, making elPrep both easier to install and use.

In addition to these new features, various performance improvements decreasing both runtime and memory use are implemented in elPrep 4, as shown by our experiments in the Benchmarks section.

### Command-line interface

The elPrep 4 software is distributed as a single binary file for Linux. A pipeline description in elPrep consists of a single command-line invocation. For example, the preparation pipeline recommended by the GATK Best Practices may look like the elPrep command shown in Listing 1.

#### Listing 1. elPrep command for executing a GATK Best Practices preparation pipeline.

~~~
elprep  sfm  input . bam  output .bam
 ––mark–duplicates  ––mark–optical–duplicates  output . metrics
 ––sorting–order  coordinate
 ––bqsr  output . recal
 ––known–sites  dbsnp_138 . hg38 . elsites
 ––bqsr–reference  hg38 . elfasta
~~~

This elPrep command executes a pipeline that takes as input a BAM file and performs (optical) duplicate marking, generates metrics, sorts the input by coordinate order, and applies base quality score recalibration, producing a single output BAM file. It is possible to specify further parameters for each option, but they are not listed here. The order in which the steps are specified is irrelevant: The elPrep implementation internally takes care of ordering the execution of the steps correctly, while also merging and parallelizing their execution. Note that the VCF and FASTA files need to be converted to an internal format beforehand, cf. the .elsites and .elfasta files in the command. These can be generated by separate elPrep commands once from the original FASTA and VCF files. The .elsites and .elfasta formats can be parsed significantly more efficiently than the VCF and FASTA formats. For more details, please consult our extensive documentation online (http://github.com/ExaScience/elprep).

### The elPrep 4 framework

elPrep, from the beginning, has been designed as a modular plug-in architecture where the implementation of SAM/BAM tools is separated from the engine that parallelizes and merges their execution [1]. While many of the core ideas from the original elPrep architecture remain unchanged, the elPrep 4 framework introduces a number of changes that make it easier to implement more complex SAM/BAM tools.

#### A phased, filtering architecture

A key idea in elPrep is to distinguish between SAM/BAM tools that can be expressed as operations on individual reads or *filters*, and operations such as sorting that operate on the whole set of reads [1]. Examples of filters include operations to remove unmapped reads, or remove reads based on genomic regions, but we have also shown that more complex operations such as duplicate marking can be expressed as filters [1].

Conceptually, elPrep distinguishes between three phases when executing pipelines:

1. Phase 1: parse the reads from file into memory while applying a first set of filters. This phase also collects all reads that are not removed by the filters into a data structure representing a SAM/BAM file;
2. Phase 2: consecutively execute all operations that use the whole set of reads. These operations can access the reads via the data structure produced in phase 1;
3. Phase 3: output the reads from memory to file while applying a final set of filters.

The elPrep 4 framework now provides hooks to extend each of these phases to execute additional operations. The main interfaces for implementing new operations are a filter interface based on higher-order functions, and the SAM data structure for representing a SAM/BAM file in memory.^2^

#### A modular plug-in architecture

The elPrep execution engine is designed as a collection of *higher-order functions* and filters that are implemented using lambda expressions [1]. Lambda expressions are anonymous, first-class functions, which allow functions to be treated as values that can be used as input values to other functions or can be used as return values. This mechanism is available in languages such as Common Lisp, C++11, Java 8, and our implementation language Go.

Concretely, elPrep models filters using two layers of filtering functions (Listing 2). The top level function receives a representation of the SAM header as an argument, so one can modify it there. This function returns another function that has a single alignment object as an argument. Code to inspect or modify an individual read goes there. The function also returns a boolean to indicate if the alignment needs to be kept in the final result output or should be removed.^3^

##### Listing 2. Skeleton structure of an elPrep filter definition.

~~~
func myFilter ( header *Header) AlignmentFilter {
       …
       **return** func ( aln *Alignment ) **bool** {
              …
              **return true or false**
       }
}
~~~

Next to the filter interface, one can also define tools that operate on the whole set of reads. The elPrep framework provides a Sam data structure that represents a SAM/BAM file in memory (Listing 3). The data structure provides access to the reads from the SAM file in the form of an array (cf. Alignments), so that whole-set operations can be expressed as parallel loops over that alignment array. We developed the Pargo [9] library for parallel programming in Go for this.

##### Listing 3. elPrep in memory representation of a SAM/BAM file.

~~~
type Sam **struct** {
        Header           *Header
        Alignments       [] * Alignment
        …
}
~~~

#### A parallel architecture

elPrep is a parallel architecture designed to take advantage of multithreading. elPrep relies on the (statically linked) Pargo library for parallel programming in Go that we developed independently [9]. The Pargo library provides various data structures for expressing parallel algorithms. Specifically, we use the following Pargo packages:

- pargo/pipeline: This package provides functions and data structures to construct and execute parallel pipelines. We use this to implement the execution of the SAM/BAM tools expressed as filters (cf. phase 1 and 3).
- pargo/sort: We use the parallel merge sort for implementing the algorithm for sorting reads by coordinate.
- pargo/sync: This package provides a parallel hash table. We use this in the implementation of various complex SAM/BAM tools such as duplicate marking, base quality score recalibration, optical duplicate marking, etc.
- pargo/parallel: This package provides various functions for parallel range-reduce operations. We use this for implementing various algorithms that operate on the whole set of reads (phase 2).

### Expressing optical duplicate marking and BQSR in elPrep 4

We added optical duplicate marking and base quality score recalibration in elPrep 4, both of which required developing new parallel algorithms that fit in the elPrep 4 framework, yet produce the same results as their counterparts in Picard/GATK 4. In the S1 Appendix, we discuss our parallel algorithm for optical duplicate marking. Similarly, in the S2 Appendix, we discuss our parallel algorithm for base quality score recalibration and application in elPrep 4.

## Results

To assess the efficiency of elPrep 4, we set up three different benchmarks where we execute a 4-step preparation pipeline specified by the GATK Best Practices [3]. We discuss raw performance by comparing the runtime and resource use of elPrep 4 versus GATK 4 and GATK 3.8. Subsequently, we discuss a scaling experiment on Amazon Web Services to compare the dollar cost of using elPrep 4 versus GATK 4.

### Benchmarks comparing elPrep 4 and GATK 4

The pipeline we benchmark contains the following steps (as specified by the GATK Best Practices [3]). We list the GATK 4 tool name for each step between brackets:

1. Sorting the BAM for coordinate order (SortSam);
2. Marking the read duplicates (MarkDuplicates);
3. Base quality score recalibration (BaseRecalibrator);
4. Applying base quality score recalibration (ApplyBQSR).

#### Data sets

We execute our benchmarks for both a whole-exome and whole-genome sequencing of NA12878. We downloaded the FASTQ files from their respective public repositories [10, 11] and aligned them using BWA mem [5]. The whole-exome sample was aligned using hg19^4^ and the whole-genome sample using hg38. The pipelines we created for both samples differ in terms of parameters used to take into account the target reference, or in case of the whole-exome sample, to use the BED file with captured regions.

#### Server and software versions

We ran our benchmarks on a 36-core server, consisting of two 18-core Intel Xeon E5-2699v3 Haswell processors clocked at 2.3GHz, allowing the simultaneous execution of up to 72 hyper-threads. The server is equipped with 256GB RAM and 2×400GB SSD disks for storing intermediate data. The machine runs Ubuntu 14.04.5 LTS. We used elPrep 4.0.0 compiled with go1.10.3, gatk-4-0.8.1 using Java 1.8.0_144, and bwa-0.7.17.

#### Whole-exome results

The benchmark results for the whole-exome data are shown in Fig. 1. There are three graphs, comparing the runtime, RAM use and disk use^5^ for GATK 4 and elPrep 4 respectively. The runtime graph shows the runtimes for each individual step in case of GATK 4 (top) versus the runtime of the merged steps in elPrep 4 for filter mode and sfm mode (bottom). The filter mode in elPrep 4 executes entirely in RAM, while the sfm mode favours disk use for intermediate results by splitting up the data by chromosomal regions for processing. The final outcomes, meaning the produced BAM, metrics and recalibration files, are the same for GATK 4 and elPrep 4 (both filter and sfm mode).

**Fig 1.**
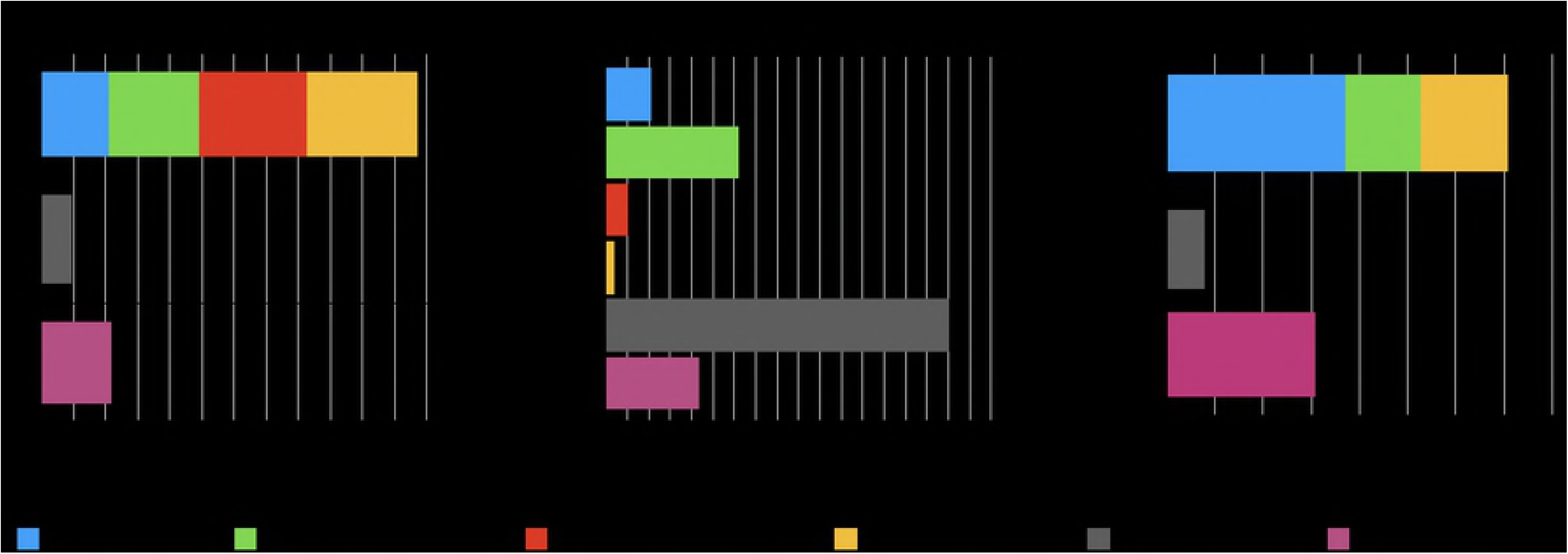
WES benchmarks. Runtime, RAM use, and disk use in GATK 4 vs. elPrep 4 (filter mode) vs. elPrep 4 (sfm mode). We see 5.4-13x speedup for 0.7-2.6x RAM use and 0.6-0.2x disk use when comparing elPrep 4 filter/sfm to GATK 4. The results, i.e. final BAM, metrics and recalibration files, are the same for all runs.

The runtime for GATK 4 is the runtimes of the individual pipeline steps added up, as the execution of these steps effectively coincide with seperate GATK 4 command-line invocations. In contrast, the results for elPrep 4 do not differentiate between the steps, as the execution of all steps is merged. The minimum RAM use of GATK 4 is determined by the peak RAM use of the individual steps, which is recorded here for the MarkDuplicates step. The minimum disk use for GATK 4 is determined by looking at the disk use of the individual steps and combining the two subsequent steps that produce the largest sum. This is a good estimate of the minimum disk space since the intermediate BAM files produced by the individual steps can be deleted once they have been processed by the next step, but not before. Here we get a peak disk use for combining the SortSam and MarkDuplicates steps.

We see that elPrep 4 (filter mode) is 13x faster, uses 2.6x more RAM, and uses only 0.15x of the disk space compared to GATK 4. Using elPrep 4 (sfm mode) we see that elPrep 4 is 5.4x faster than GATK 4, using only 0.7x the RAM and 0.6x the peak disk space that GATK 4 uses. Concretely, we go from a runtime of 58m31s using 31GB of RAM and 26.34GB of disk in GATK 4 to a runtime of 4m35s using 80GB RAM and 4GB of disk for the elPrep 4 filter mode, or a runtime of 10m57s using 22GB RAM and 15.5GB of disk for the elPrep 4 sfm mode.

Overall, elPrep 4 executes the pipeline faster, while making more efficient use of the compute resources (RAM/disk/threads) than GATK 4, in both filter and sfm modes.

#### Whole-genome results

The results for our whole-genome benchmark are shown in Fig. 2, comparing runtimes, RAM use and disk use for GATK 4 and elPrep 4 (sfm mode). We see that elPrep 4 executes the pipeline 7.4x faster than GATK 4, while using 0.84x of the RAM and just 0.7x of the disk space. The runtime goes down from almost 27h in GATK 4 to roughly 3h37m in elPrep 4, while RAM use goes down from roughly 229GB in GATK 4 to 192GB in elPrep 4, and the peak disk use goes down from 520GB in GATK 4 to 346GB in elPrep 4. Again, elPrep 4 achieves these speedups while producing the same results compared to the GATK 4 run.

**Fig 2.**
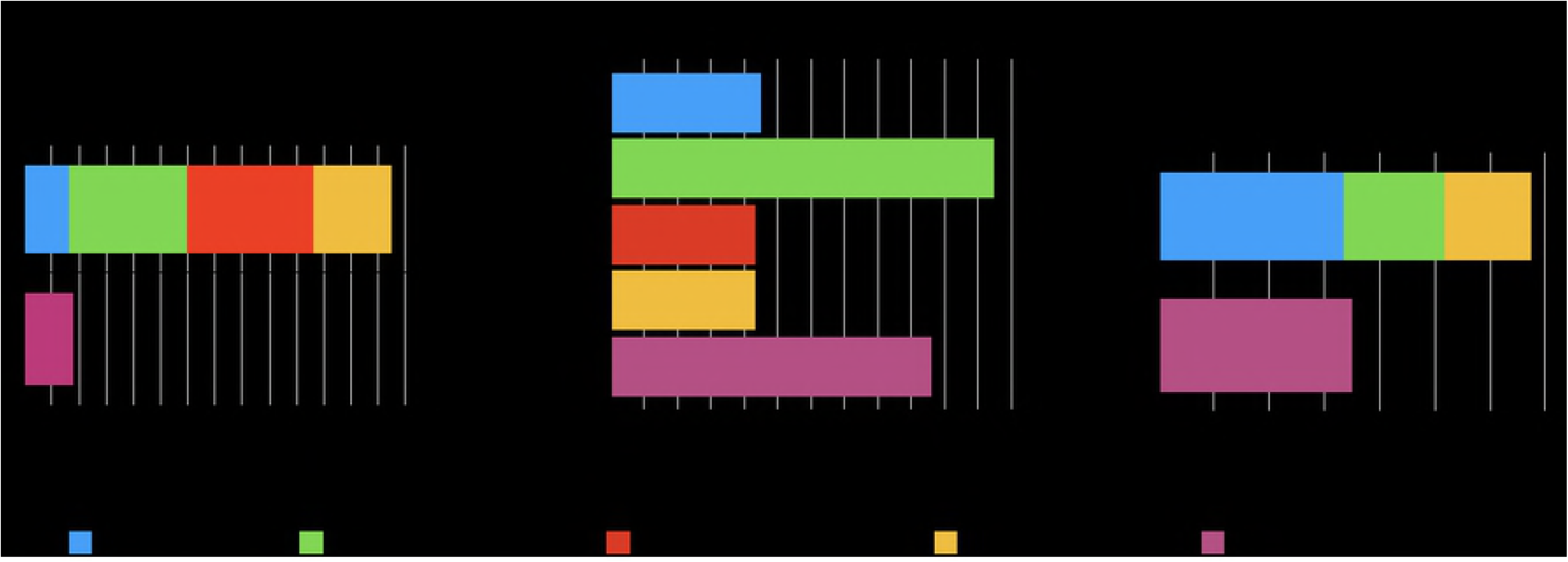
WGS benchmarks. Runtime, RAM use, and disk use in GATK 4 vs. elPrep 4 (sfm mode). elPrep 4 executes the pipeline 7.4x faster than GATK 4, using 0.84x of the RAM, and only 0.7x of the disk space. The final BAM, metrics, and recalibration files are the same for both runs.

#### Comparison of outputs elPrep 4 and GATK 4

elPrep 4 produces the same output as GATK 4. When we reimplement a tool from GATK 4, Picard, or SAMtools, our goal is to come up with a new algorithm that takes advantage of elPrep’s parallel architecture, yet does not change the semantics of the original algorithm. This means that we try to respect the heuristics, execution order, etc. of the original algorithms as much as possible, so that the outcomes are the same.

One challenge is that many of the algorithms are non-deterministic. For example, the GATK 4/Picard mark duplicate algorithm compares reads for duplicate marking by comparing the adapted mapping positions and adapted quality scores. When two reads have the same adapted mapping position, the idea is to mark the read with the worse adapted quality score as a duplicate. It may however occur that two reads have the exact same mapping position *and* the exact same quality score. In this case, which read is marked as the duplicate, conceptually does not matter, and in Picard and GATK 4, which one is marked will just depend on the order of the reads in the input file. Since elPrep parallelizes the processing of reads, they are not always examined in the same order of the input file. Because of this, there may be small differences when comparing BAMs, albeit not meaningful differences. In previous work we discussed how to run elPrep in a deterministic mode for duplicate marking to compare BAMs between GATK 4/Picard and elPrep exactly using Unix diff [1]. One can now in addition compare the metrics files that are generated with optical duplicate marking using Unix diff or MultiQC.

Similarly, we can show that the base quality score recalibration (BQSR) algorithm in elPrep 4 produces the exact same result as GATK 4. We can verify this by comparing the .recal files that contain the BQSR statistics and are generated by both tools using Unix diff or MultiQC. The BQSR algorithm takes into account duplicated reads for calculating these statistics, and since duplicate marking is non-deterministic, an exact comparison between GATK 4 and elPrep 4 only makes sense when they are passed the exact same input BAM for BQSR calculation. So when we call GATK 4 and elPrep 4 with a BAM file that is already coordinate sorted and marked for duplicates, we see that the .recal files that are produced by both tools when performing BQSR are exactly the same when doing a Unix diff command. We can also compare the BAMs produced by GATK 4 and elPrep 4 using Unix diff, but it is important to first sort the optional fields in each read, and sort the files using Unix sort. The latter are needed to handle the non-deterministic order of the optional fields on the one hand (see SAM/BAM specification [2]), and the non-determinism of sorting for coordinate order –when multiple reads have the same mapping positions. A recipe for comparing the execution of GATK 4 and elPrep 4 is given below:

1. Sort input BAM by query name to handle non-determinism of the coordinate sort in the next step;
2. Sort + mark the input BAM for duplicates (using elPrep or GATK/Picard);
3. Run elPrep with –bqsr and –deterministic mode on the BAM from step 2;
4. Run GATK with BaseRecalibrator and ApplyBQSR on the BAM from step 2;
5. Perform a Unix diff on .recal files created by elPrep and GATK runs;
6. Remove PG tag and sort optional fields of elPrep and GATK output BAMs (using biobambam [14]);
7. Unix sort elPrep and GATK SAMs;
8. Perform Unix diff on elPrep and GATK SAMs.

The restrictions that are needed for introducing determinism in the pipeline executions for exact comparisons are in general not recommended when using elPrep 4. They create performance bottlenecks without providing any interesting additional information, and are only useful for verifying elPrep 4’s equivalence to GATK 4.

### Benchmarks comparing elPrep 4 and GATK 3.8

The pipeline we benchmark for comparing the performance of elPrep 4 and GATK 3.8 is the same pipeline as the one used for the comparison with GATK 4, but the difference is that Picard tools are used for some of the steps. The functionality of Picard tools and GATK is merged in GATK 4, but for earlier versions of GATK, Picard tools is the standard tool for implementing some of the pipeline steps [3].

Below we list the pipeline steps and the tool that is recommended for processing them in the GATK Best Practices [3] for GATK versions predating GATK 4:

1. Sorting the BAM for coordinate order (SortSam from Picard);
2. Marking the read duplicates (MarkDuplicates from Picard);
3. Base quality score recalibration (BaseRecalibrator from GATK);
4. Applying base quality score recalibration (PrintReads from GATK).

#### Data sets

We benchmark the same whole-genome data set that we use in our benchmarks for GATK 4, namely the Illumina Platinum whole-genome sequencing of NA12878 [11]. We created the aligned BAM file from the original FASTQ files by aligning the data against hg38 using bwa mem.

#### Server and software versions

We ran our benchmarks on the same 36-core server we use for our GATK 4 benchmarks. We used elPrep 4.0.0 compiled with go1.10.3, gatk-3.8.0 using Java 1.8.0_144, picard-tools-2.9.2, and bwa-0.7.17.

#### Whole-genome results

The benchmark results comparing GATK 3.8 and elPrep 4 are shown in Fig. 3. They compare runtime, RAM, and disk use. elPrep 4 executes the pipeline more than 18x faster than GATK 3.8, while using only 0.85x of the peak RAM and 0.8x of the peak disk space that GATK 3.8 uses. Concretely, the runtime goes down from almost 65h to roughly 3h40m, while peak RAM use goes down from 225GB to 192GB, and peak disk use from 442GB to 350GB. Note that the total runtime for GATK 3.8 is the sum of the runtimes of the individual steps. The peak RAM use for GATK 3.8 is the largest RAM use of the individual steps. The peak disk use of the GATK 3.8 run is calculated as the sum of the disk use for the SortSAM and MarkDuplicates steps. For elPrep 4, all of the pipeline steps are merged and consequently so are the results presented in the figures.

**Fig 3.**
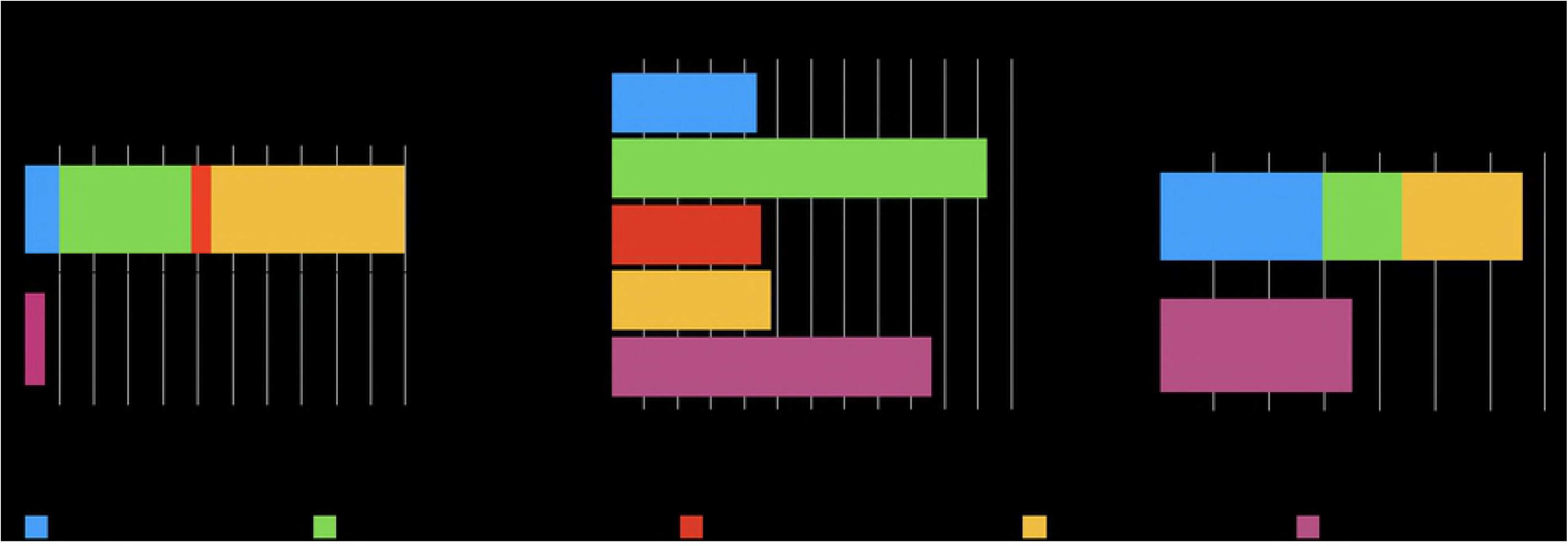
WGS benchmarks. Runtime, RAM use, and disk use in GATK 3.8 vs. elPrep 4 (sfm mode). elPrep 4 executes the pipeline 18.2x faster than GATK 3.8, using 0.85x of the RAM, and only 0.8x of the disk space.

Note that we only compare the raw performance of elPrep 4 and GATK 3.8. The algorithms and outcome of the BQSR tools in GATK 4 changed compared to GATK 3.8. Since elPrep 4 implements the GATK 4 algorithm, an exact comparison of outcomes between elPrep 4 and GATK 3.8 is not possible, as is the case when comparing the outcomes of GATK 4 and GATK 3.8.

### Scaling experiment on Amazon Web Services

We set up a scaling experiment on Amazon Web Services (AWS) cloud servers (EC2) that uses the same 4-step pipeline (sorting, duplicate marking, base quality score recalibration and application) that is used for comparing the raw performance of GATK 4 and elPrep 4 in the previous sections. In this experiment, we measure the runtime on a wide range of EC2 instances with different numbers of CPUs and amounts of RAM, which allows us to assess the scaling behavior of GATK 4 and elPrep 4. We also calculate the cost of running the benchmark on each instance based on Amazon EC2 on-demand pricing. We show that elPrep scales better and therefore has a stable cost across different configurations, whereas the cost to speed up GATK 4 by allocating more compute resources increases rapidly.

#### Data sets

We use the same whole-exome and whole-genome data sets that we use in the rest of the benchmarks for comparing GATK 4 and elPrep 4. Hence, we use the genome-in-a-bottle whole-exome for NA12878 aligned against hg19 [10], and the Illumina Platinum whole-genome for NA12878 aligned against hg38 [11].

#### Server and software versions

We ran the pipeline for both whole-exome and whole-genome data sets on a wide range of Amazon instances, as listed in Table 1. The table lists the name of the instance, followed by the number of virtual CPUs, the amount of virtual RAM, and the dollar cost per hour^6^ to rent such an instance. All of the Amazon instances run Amazon Linux 2. We additionally installed elPrep 4.0.0 compiled with go1.10.3, gatk-4-0.8.1 using Java 1.8.0_144, and bwa-0.7.17 for running the benchmarks.

**Table 1.**
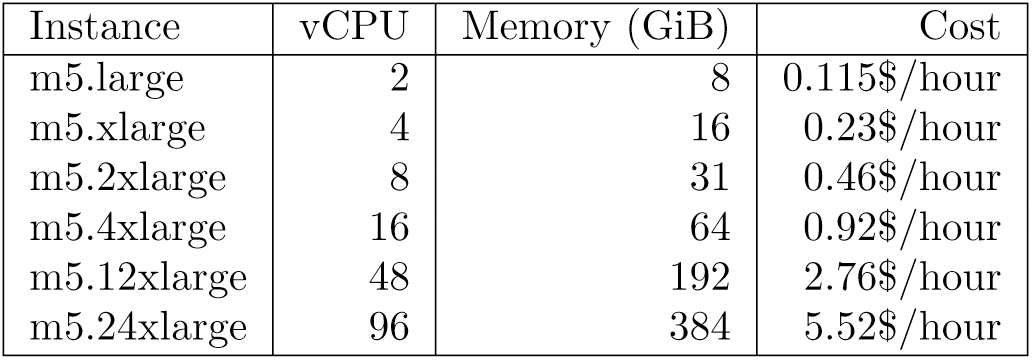
AWS instances used in our benchmarks. Prices for EU (Frankfurt) Oct. 2018.

#### Whole-exome results

The results for running our whole-exome benchmark on AWS are shown in Fig. 4. The figure shows both the dollar cost and runtime for comparing the GATK 4 and elPrep 4 runs on Amazon instances ranging from m5.large to m5.24xlarge. The dollar cost is calculated per run by multiplying its runtime by the dollar cost per hour^7^ for each Amazon instance type, as listed in Table 1. While the GATK 4 runtime scales somewhat with using a larger instance, the scaling for elPrep 4 is much better, as the runtime is nearly halved with each instance increase. The dollar cost goes up steeply for GATK 4 with each instance increase. In contrast, because elPrep 4 scales so well with the increase of compute resources, the dollar cost per run only increases slightly for each instance increase.

**Fig 4.**
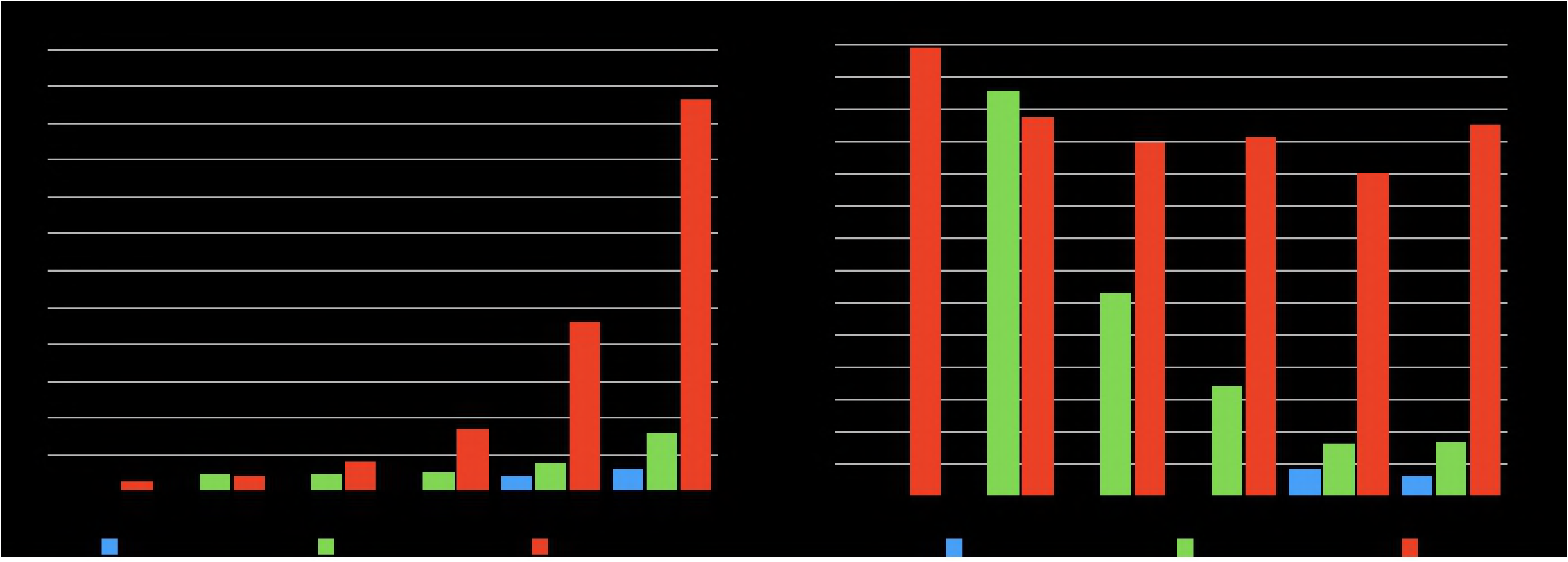
AWS WES benchmarks. The dollar cost and runtime on Amazon Web Services for running a 4-step pipeline on a whole exome using GATK 4 versus elPrep 4 (filter and sfm modes). The runtime of elPrep 4 scales linearly with the increase of compute resources, while GATK 4 shows only limited improvements. The dollar cost per run increases steeply with GATK 4 for little performance improvements, while the dollar cost with elPrep 4 remains mostly stable across all Amazon instances.

The cheapest run of the whole exome is observed for GATK 4 on instance m5.large, where it runs for 69m34s for 0.13$. The cheapest run with elPrep 4 is on instance m5.2xlarge with a runtime of 31m38s for 0.24$ using the elPrep sfm mode. This means the cheapest elPrep 4 run is roughly 2x faster for roughly 2x the cost of the cheapest GATK 4 run. The fastest run of the benchmark is with elPrep filter mode on instance m5.24xlarge, taking 3m25s and costing 0.31$. The fastest run with GATK 4 uses instance m5.12xlarge and takes 50m6s, costing 2.30$. Hence the fastest elPrep 4 run is almost 15x faster than the fastest GATK 4 run, and costs 7.5x less.

#### Whole-genome results

The AWS benchmark results for our whole-genome sample are shown in Fig. 5. Both the dollar cost and runtime for GATK 4 and elPrep 4 runs are shown for different Amazon instances. The elPrep 4 benchmark was only run on instance m5.24xlarge, because it is the only instance that satisfies the elPrep memory requirements for this particular whole-genome data set. In contrast, the GATK 4 runs are able to execute on Amazon instances ranging from m5.large to m5.24xlarge.

**Fig 5.**
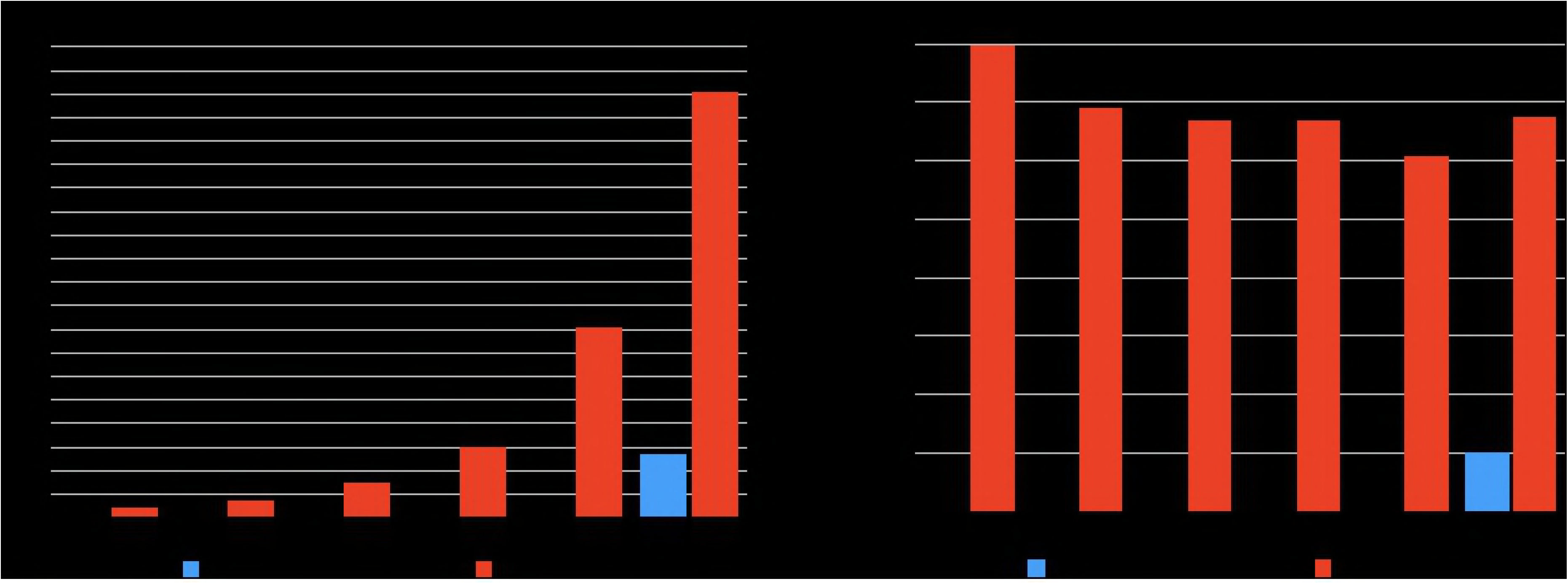
AWS WGS benchmarks. The dollar cost and runtime on Amazon Web Services for running a 4-step pipeline on a whole genome using GATK 4 versus elPrep 4. While GATK 4 is able to run on a wider range of Amazon instances, the overall runtime is much larger compared to elPrep 4. The fastest run with GATK 4 takes over 17.5 hours on m5.12xlarge and costs 48.71$, whereas the elPrep 4 run takes a bit less than 3 hours and costs only 16.25$ on m5.24xlarge, being almost 6x faster for 3x less money.

Similar to the whole-exome results, the overall cheapest run is for GATK 4 on m5.large, costing 2.68$, but taking 23h17m. The elPrep 4 run on m5.24xlarge costs 16.25$, but only takes 2h57s. So the elPrep 4 run is almost 8x faster and costs only 6x more. The fastest GATK 4 run is recorded on instance m5.12x large and takes 17h39m at a cost of 48.71$. This means the elPrep 4 run is almost 6x faster and 3x cheaper than the fastest GATK 4 run.

## Related Work

There is a large body of related work to speed up DNA sequencing pipelines. First of all, the GATK 4 team at Broad Institute is also developing an alternative implementation of GATK 4 in Spark [12]. While GATK 3.8 and earlier versions had options for configuring multithreading, these are mostly removed from the standard GATK 4 implementation.^8^ Instead, the idea is to use the GATK 4 Spark implementation in place of GATK 4 for coarse-grained parallelization. Whereas elPrep focuses on single-node optimizations through multithreaded programming, Spark is optimized for parallelization on a compute cluster [12]. The GATK 4 Spark implementation is currently only available as a beta release, and initial tests show results that differ from the reference GATK 4 implementation, making it difficult to compare to elPrep. Also, the general strategy behind the GATK 4 Spark implementation is to parallelize the individual Spark GATK 4 tools, whereas elPrep combines and merges the execution of several tools, which we have shown to be more scalable and efficient [1].

Similarly, there are many tools such as bamUtil [13], biobambam [14], and Sambamba [15] that focus on optimizing individual pipeline steps, but do not combine the execution of multiple steps, overall yielding a worse performance than elPrep or producing different results [1]. A more recent approach is Sentieon, which promises a 10-fold speedup compared to GATK variant calling while producing identical results [16]. They offer a reimplementation of the GATK 3.5 variant caller that is optimized for multithreading, but this implementation is closed source.

We previously discussed related work that focuses on optimizing the whole sequencing pipeline by stepping away from community-defined standards such as the SAM/BAM format to define their own data formats and new algorithms for processing them [1]. Examples we previously discussed [1] include ISAAC [17] and BALSA [18] for GPUs, and more recent approaches such as Dragen [19] and Genalice [20] that promise considerable speedups compared to standard tools. Both Dragen and Genalice are commercial tools that implement their own patented algorithms for implementing a full variant calling pipeline. The outcomes therefore differ from the community-defined reference pipelines such those based on the GATK Best Practices. Dragen additionally requires specialized hardware in the form of FPGAs to run. In contrast, elPrep is an open-source implementation that focuses on supporting the community-based standards such as SAM/BAM/VCF/BED, offers the flexibility to configure the pipelines, and targets multicore servers as generally available in, for example, cloud services.

## Conclusions

elPrep 4 is a reimplementation of the elPrep framework [1] for processing sequence alignment map files (SAM/BAM) in the Go programming language. It introduces new and improved functionality for sorting, optical duplicate marking, base quality score recalibration, MultiQC-compatible metrics, and various filtering options. This allows elPrep to process most of the preparation pipelines defined by the GATK Best Practices [3], but also other types of pipelines [7]. For this, we developed new parallel algorithms that reimplement the GATK 4 tools for optical duplicate marking and base quality score recalibration in the elPrep 4 framework, greatly speeding up the execution of these steps compared to GATK 4, while producing the same results.

In our benchmarks, we compare the raw performance of elPrep 4 to GATK 4 and GATK 3.8, on both a whole-exome and whole-genome data sample of NA12878 (Genome in a bottle/Illumina Platinum genome). Compared to GATK 4, elPrep 4 executes a 4-step pipeline consisting of sorting, duplicate marking, base quality score recalibration and application, 7.4x faster, while using less RAM and disk space. Similarly, elPrep 4 executes the same pipeline more than 18x faster than GATK 3.8, using fewer RAM and disk resources. We ran a scaling experiment on Amazon Web Services (AWS) to compare the runtime and dollar costs of running the 4-step pipeline on a wide range of Amazon compute instances using elPrep 4 and GATK 4. elPrep 4 makes better use of the available compute resources such as CPUs and RAM than GATK 4. The cost of using elPrep 4 on AWS more or less remains stable when using a more expensive AWS instance because of the good scaling. Concretely, the fastest elPrep 4 run of the 4-step pipeline on WES data is 15x faster (3m25s vs 50m6s) and 7.5x cheaper (0.31$ vs. 2.30$) than the fastest GATK 4 run. The overall cheapest run is for GATK 4, costing 0.13$, but also taking around 70m. Similarly, the fastest elPrep 4 run on AWS for WGS data is 6x faster (less than 3 hours versus 17.5 hours) than the fastest GATK 4 run, costing 3x less (16.25$ vs. 48.71$). Again, overall the cheapest run is recorded for GATK 4 at 2.68$, but it then takes almost 24 hours.

elPrep 4 differs from related work in its approach to optimizing sequencing pipelines. Rather than optimizing individual tools, the elPrep 4 framework executes a pipeline by defining an optimal ordering of the steps, and merges and parallelizes their execution, which overall yields a better speedup. elPrep 4 achieves its speedups while offering the flexibility to freely plug pipeline steps in or out, and producing the same results as reference implementations of these steps in GATK 4, Picard, and SAMtools. elPrep 4 works with community-defined standards such as SAM/BAM/VCF/BED rather than defining its own formats for achieving its speedups, making elPrep 4 (backwards) compatible with other standard tools and workflows [7, 21, 22].

## Supporting information

**S1 Appendix. Expressing optical duplicate marking in elPrep 4.** We describe how to express the optical duplicate marking algorithm from Picard/GATK 4 as a parallel, single-pass algorithm in the new elPrep 4 framework.

**S2 Appendix. Expressing base quality score recalibration (BQSR) in elPrep 4.** We explain how to express the base quality score recalibration and application algorithms (BQSR) from GATK 4 as a parallel, map-reduce algorithm in the new elPrep 4 framework.

## Footnotes

1 The recent release of GATK 4 contains preparation tools subsuming the Picard software.

2 The original elPrep framework only makes it easy to add new filter operations. Sorting was the only whole-set operation, and its implementation was integrated with the elPrep framework.

3 The original elPrep interface for defining filters in the Common Lisp implementation had three layers of functions. In between the header and alignment filter, there was a function for thread-local storage, but this works differently in Go.

4 We use hg19 for the genome-in-a-bottle whole-exome sample so that we can use the hg19-compatible BED file with captured regions that comes with the sample.

5 Disk use refers to the number of GBs written to disk while executing the pipeline steps.

6 On-demand pricing in EU (Frankfurt) as of October 2018.

7 In practice, on AWS, the cost is rounded up for each hour started, but we did no rounding in our calculations.

8 GATK 4 still relies on multithreading for libraries that implement compute-intensive kernels (e.g. PairHMM), as well as the multithreading used by the JVM (e.g. for garbage collection).

